# The $10 proteome: low-cost, deep and quantitative proteome profiling of limited sample amounts using the Orbitrap Astral and timsTOF Ultra 2 mass spectrometers

**DOI:** 10.1101/2025.07.29.667408

**Authors:** Siqi Huang, Chao Wang, Hsien-Jung L. Lin, Ryan T. Kelly

**Affiliations:** Department of Chemistry and Biochemistry, Brigham Young University, Provo, UT 84602

## Abstract

Mass spectrometry (MS)-based proteomics remains technically demanding and prohibitively expensive for many large-scale or routine applications, with per-sample costs of hundreds of dollars or more. To democratize access to proteomics and facilitate its integration into more high-throughput multi-omic studies, we present a robust analytical framework for achieving in-depth, quantitative proteome profiling at a cost of approximately $10 per sample, termed the “$10 proteome.” Using the Thermo Fisher Orbitrap Astral and Bruker timsTOF Ultra 2 mass spectrometers, we evaluated performance across sample inputs ranging from 200 pg to 100 ng and active gradient lengths from 5 to 60 minutes. Proteome coverage saturated within the low-nanogram input range, with ∼8000 protein groups quantified from as little as 10 ng of input and nearly 6000 protein groups from 200 pg. With already demonstrated low-cost one-pot sample preparation workflows that are appropriate for this sample input range, standardized MS acquisition settings, and high-throughput nanoLC operated at ∼10 min per sample, the $10 proteome becomes feasible. This study establishes a practical, scalable, and cost-effective foundation for global proteome profiling, paving the way for routine, large-scale applications in systems biology, clinical research and beyond.

## Introduction

Over the past two decades, the cost of DNA and RNA sequencing has plummeted, dramatically transforming the landscape of genomics and transcriptomics research. The price of sequencing a human genome has dropped from roughly $100 million in 2001 to less than $1,000 as of 2024 (1), a trend driven by technological innovations in next-generation sequencing, improvements in throughput, and extensive commercialization. This democratization of sequencing technologies has enabled widespread applications in research and clinical settings, including precision oncology, population-scale studies, and rapid pathogen detection (2-5). The efficiency and affordability of sequencing have also spurred the development of large consortia and biobanks, generating massive multiomic datasets that continue to reshape biomedical research, diagnosis and treatment (6-8).

In contrast to nucleic acid sequencing, proteomics, the large-scale study of proteins and their functions (9-11), has not benefited from similar cost reductions or scalability, despite its essential role in understanding functional biology. Mass spectrometry (MS)-based proteomics, the cornerstone of high-resolution protein analysis, remains technically demanding and resource-intensive (12-14). Per-sample costs can range from $200 to over $1,000, depending on the depth of proteome coverage, sample complexity, and the specific instrumentation used (15-17). As a result, MS-based proteomics remains prohibitively expensive for applications like routine screening.

Several factors contribute to the high cost. Liquid chromatography (LC) and MS instruments are expensive, depreciate quickly and require service contracts or unpredictable maintenance costs. Standardization of workflows and instrumentation could significantly reduce expenses (18, 19), yet the diversity of biological samples, including suspended cells, biofluids, solid tissues, complex microbial communities, etc. poses a significant challenge to this end. Sample amounts can also vary from subcellular to bulk scale, and preparation protocols currently must be tailored accordingly, requiring substantial technical expertise. Prepared samples, with protein quantities ranging from picograms to tens of micrograms, are then analyzed by LC-MS. The vast array of options, including LC columns, flow rates, mass spectrometers, acquisition settings, and data analysis strategies, further complicates standardization. These diverse inputs and analytical variables hinder the development of streamlined, cost-saving workflows, unlike the more unified approaches seen in nucleic acid sequencing. Additionally, continuous advances in LC-MS technologies and data acquisition methods, such as the rapid shift from data-dependent to data-independent acquisition (DIA) for label-free analysis, have improved proteome coverage and data completeness (20, 21), but further delayed methodological convergence.

Due to these challenges, proteomics remains a relatively small enterprise, typically confined to specialized laboratories and reliant on significant technical expertise and infrastructure (12, 22). Smaller research labs and core facilities often struggle to accommodate the diversity of incoming samples, leading to a low success rate for sample analysis. These limitations have hindered the widespread adoption of proteomics in large-scale studies and clinical workflows, preventing it from achieving the transformative impact seen with genomics, even though proteins execute the majority of cellular functions.

Closing this gap requires the development of standardized, automated, and low-cost LC-MS workflows. As we establish here, recent advances now enable in-depth, quantitative global proteome profiles to be obtained for as little as $10 per sample. This can be achieved by:

- Implementing simplified sample preparation methods that were originally developed for single-cell and spatial proteomics but that are equally effective for inputs of hundreds of cells (up to ∼50 ng of protein)
- Employing a limited menu of LC-MS acquisition settings across all samples, eliminating the need for time-consuming parameter optimization for every study
- Utilizing the latest generation of high-performance mass spectrometers (e.g., Bruker timsTOF SCP/Ultra/Ultra 2/AIP or Thermo Fisher Orbitrap Astral/Astral Zoom) with DIA for significantly improved speed and sensitivity
- Limiting run times to ∼10 minutes per sample to maximize the productivity of the expensive mass spectrometers
- Automating data processing, analysis, summarization, and visualization to streamline the end-to-end workflow

In the present study, we present key advances toward achieving the $10 proteome. Using both the Orbitrap Astral and timsTOF Ultra 2 mass spectrometers, we first evaluated sample inputs ranging from 0.2 ng (single-cell equivalent) to 10 ng of total protein. For both instruments, proteome coverage began to plateau at the higher end of this range, indicating speed-limited saturation. Notably, this optimal input range aligns with recently developed one-pot (23-29) and/or one-step (30, 31) sample preparation strategies, which enable the preparation of thousands of samples per day at a cost of just a few cents each. We employed fixed MS acquisition settings on both the timsTOF Ultra 2 and Orbitrap Astral systems across all sample inputs and gradient lengths, achieving strong and consistent performance without the need for extensive parameter optimization. Proteome coverage for low-nanogram inputs approached saturation with 5–20-minute gradient lengths while maintaining high reproducibility and linearity between input amount and quantified protein abundances. We then analyzed samples ranging from 0.2 to 100 ng using a dual-column nanoLC system operating at 126 samples per day (∼11.4 minutes per sample), approaching the target of 10 minutes per run. Under these conditions and when performing data analysis using Spectronaut 19 software, we achieved an average proteome coverage of ∼6,000 protein groups for 200 pg and ∼8,000 protein groups for 20–100 ng inputs. When combined with automated data analysis using available search tools and our open-source MSConnect data management platform (32), these developments make robust, in-depth global proteome profiling of nearly any sample type feasible for approximately $10 in direct costs per sample.

## Results and Discussion

We sought to demonstrate that reproducible, quantitative and in-depth proteome profiling can be achieved at ultra-low cost using the flagship Orbitrap Astral and timsTOF Ultra 2 instruments from Thermo Fisher Scientific and Bruker Corp., respectively. One-pot (24, 27, 28) (and more recently one-step (30, 31)) sample preparation has been demonstrated across the range of sample inputs tested here. This has included one to hundreds of cells as well as fresh frozen and formalin-fixed, paraffin embedded tissues of various sizes (25, 26, 28, 30, 31, 33-44). For these small samples, the cost of reagents is just a few cents, and the sample amount is sufficiently small that online cleanup is sufficient. That is, expensive offline cleanup kits become unnecessary. Here, we sought to demonstrate that such samples could also be analyzed at low cost. The *analytical* (LC-MS) strategy to achieve a $10 proteome, which is the focus of the present study, combines the following elements: (1) Using a single or limited number of MS acquisition methods across all sample inputs and analysis times, thus avoiding costly method development. (2) Limiting the amount of sample prepared and analyzed to the low-nanogram range regardless of sample availability to make use of the ultra-low-cost sample preparation strategies discussed above. (3) Leveraging the fast acquisition rates of new mass spectrometers to achieve in-depth proteome profiles with short gradients.

We note that since these experiments were performed, Thermo Fisher Scientific has released the Orbitrap Astral Zoom mass spectrometer (45, 46), and Bruker Corp. has released the timsUltra AIP (47). As such, the instruments tested are no longer the latest offerings from either vendor. With newer generations of instruments and further advances in separations and search algorithms, the proteome depth and/or cost per sample will be further improved.

### Cost analysis to achieve a $10 proteome

For the purposes of our calculations, we assume that the Orbitrap Astral and timsTOF Ultra series mass spectrometers have a street price of $1,000,000 USD and a service contract price of $60,000/year for Years 2–8. The instruments are assumed to fully depreciate within 8 years. An $80,000 LC system depreciates over the same period and carries a service contract of $7,000/year for Years 2–8. This results in an instrument cost of $530/day. We allocate $120,000/year for salaries plus 23.7% for benefits, which is the current benefits rate at our university. This could cover two salaried technicians, each earning $60,000/year, or a more highly skilled employee earning a larger salary plus a part-time assistant, etc. The cost of consumables (reagents, solvents, well plates, LC columns, pipette tips, etc.) is covered by $215/day. Note that trypsin/lys-C can be a significant expense for standard high-input proteomics, but these reagents will cost no more than $0.04/sample since we will use less than 10 ng/sample and the price is of the Rapid Trypsin/Lys-C kit we used is $400/100 µg. Also, the cost of sample cleanup kits (e.g., S-Trap) is unnecessary for the sub-50 ng samples being analyzed. Data management, results summarization and visualization can be automatically performed using open-source software such as MSConnect (32). This amounts to direct costs of ∼$1150/day to cover instrumentation, reagents and labor associated with preparing samples for and operating a single mass spectrometer.

A 10-minute analysis time equates to 144 samples per day, yet we conservatively assume that instruments will collect data from samples 80% of the time, while the other 20% of the time is devoted to maintenance, repair and analysis of quality control samples. This leaves an average of 115 samples analyzed per day, or $10/sample. A multi-column nanoLC system with a ∼100% duty cycle, as demonstrated below, may be employed such that the active elution time matches the total cycle time.

### A standardized acquisition method and ≤10 ng input enable low-cost, deep proteome profiling

In MS-based proteomics, factors such as sample input, LC gradient length, column properties, flow rate, mass spectrometer model and acquisition parameters interact to influence performance, and fully optimizing all variables would require an impractically large experimental matrix. Instead, we adopted a single, broadly effective set of acquisition parameters that balance performance across conditions. Thanks to the substantial advances in modern mass spectrometers, even unoptimized settings yield markedly improved proteome coverage in significantly less time compared to just a few years ago. We assessed performance across five sample inputs (0.2–10 ng), six gradient lengths (5–60 min), and two mass spectrometers (timsTOF Ultra 2 and Orbitrap Astral), keeping the column and acquisition settings constant throughout for each mass spectrometer. Avoiding extensive parameter optimization allowed us to generate data within a few days rather than months. We propose that such a ‘one-size-fits-most’ strategy is critical for standardizing workflows and reducing the cost of high-throughput proteome profiling.

The selected MS acquisition method for each mass spectrometer was first evaluated across sample loadings in the range of 0.2 ng to 10 ng, roughly equivalent to 1–50 cells, while holding the active gradient constant at 20 min. As shown in Figure 1A, proteome coverage was quite similar for both the Orbitrap Astral and the timsTOF Ultra II mass spectrometers across this range of sample inputs, with the Astral providing slightly greater proteome coverage for 0.2 ng input and the timsTOF quantifying ∼8000 protein groups for 10-ng samples. Reproducibility was also robust for both instruments under these conditions, with median coefficients of variation (CVs) as low as 7.3% at 10 ng sample input and 16.6% at the single-cell level (200 pg), as shown in Figure 1B. Notably, a fivefold increase in sample input from 200 pg to 1 ng produces a substantial 44% increase in proteome coverage for the Orbitrap Astral, while another fivefold increase in sample size from 2 to 10 ng increased proteome coverage by just ∼7%. The respective increases for the timsTOF Ultra 2 were 50% and 14%. Together, these results indicate that the instruments are approaching their speed-limited saturation points, and we might expect minimal further improvement beyond the 10-ng input range. These findings are also consistent with those of both instrument vendors. For example, in a dilution series analyzing 62.5 pg to 16.0 ng using the timsTOF Ultra 2 mass spectrometer, researchers from Bruker Corp. showed clear evidence of saturation, with a doubling in sample input from 8 to 16 ng only increasing proteome coverage by 0.7% or 7% depending on how the search was performed (48). Note, the proteome coverage we report here using the same instrument model is greater at 10 ng than the authors reported for 16 ng despite our use of a shorter active gradient. This is likely due to our lower flow, more sensitive separations. Similarly, researchers from Thermo Fisher Scientific showed that proteome coverage for a 3-species mixture only increased by 1% when sample input doubled from 10 to 20 ng on the Orbitrap Astral with FAIMS Pro interface, again pointing to saturation in the low-nanogram range for these modern mass spectrometers (49).

**Figure 1.**
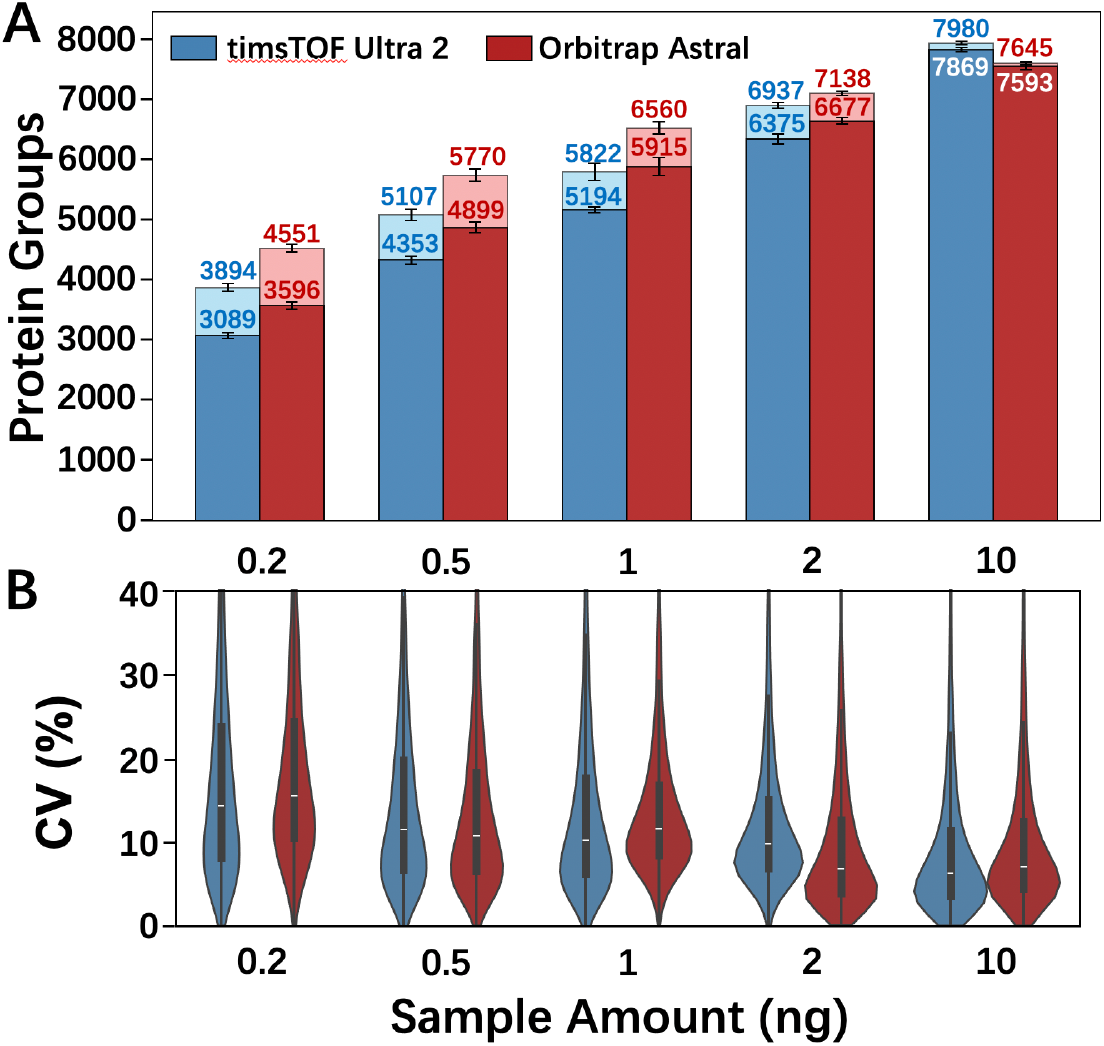
Varied sample amount with fixed MS acquisition methods and gradient length (20 min). (A) Proteome coverage. Lighter shading indicates additional proteins identified using Match Between Runs in DIA-NN. (B) Coefficients of variation.

These results have important implications for bulk-scale proteome profiling. For many studies, samples containing a microgram or more of protein may be available. However, preparing such large quantities and then analyzing only ∼10 ng is highly wasteful. For example, it can be more difficult to extract proteins from larger samples considering their smaller surface area-to-volume ratios, which can then necessitate the use of harsh surfactants (e.g., SDS) that must be removed later in the workflow. This cleanup adds to both labor and material costs. In addition, large samples require more costly protease. Alternatively, if only ∼10 ng of sample is prepared in the first place, one-pot sample preparation becomes feasible. These one-pot preparation protocols such as nanoPOTS (24), microPOTS (27) and autoPOTS (28) have proven equally effective for samples containing one to hundreds of cells. We have since simplified our one-pot protocol to a single reagent addition step following incubation for 1 h using mere pennies of protease per sample and requiring no offline sample cleanup (30, 31). Thus, preparing only what is needed for the final proteomic analysis can dramatically reduce costs. When samples comprise primary or cultured cells, microdissected tissues or biofluids (e.g., plasma), it should be relatively straightforward to isolate only the desired amount of starting material by respectively cell counting, limiting the dissected areas, or, in the case of biofluids, relying on known protein concentrations. For other samples such as biopsied tissues, protein extraction followed by a BCA assay may be required prior to aliquoting and digesting only the desired nanogram sample. This may add some additional complexity and cost to the workflow but will still result in saved protease and cleaner LC columns, etc. Up-front protein isolation and quantification is also required for popular modern bulk-scale proteomics sample preparation approaches. For example, after ensuring that 1–100 µg of material has been isolated and assayed for total protein content, the S-Trap kit and protocol then requires 20 discrete steps using 11 different reagents (50). Clearly, simple sample preparation geared toward low-input analysis will result in significant cost savings and is essential for achieving a $10 proteome.

### Label-free quantification for low-input samples

We investigated peptide peak intensities as a function of sample amount. The MS1 peak intensity increased almost perfectly linearly with sample input on the Orbitrap Astral, with an average coefficient of determination (R-squared value) of 0.996 for peptides that were detected in all sample loadings (Figure 2A). Thus, there was no indication of saturation throughout the ion path. The measured signal intensities vs. sample input was also robust but less linear for the timsTOF Ultra 2, with an average R-squared value of 0.945. When the 10-ng samples were excluded from the analysis, the mean R-squared value increased to 0.985 (Figure 2A), which points to some saturation even at the low input level of 10 ng. We investigated the source of this nonlinearity by binning the detected peptides into 5 groups according to their observed linearity between sample input and signal and plotting the distribution of ion mobilities for each quantile. As shown in Figure 2B, the peptides exhibiting the poorest linearity of signal vs sample amount have the highest average mobility (lowest 1/k_0_ value). This points to a space-charge limitation, likely in the tims cartridge, as higher mobility ions are ejected at a greater rate when space charge effects become significant (51). As such, the somewhat limited linearity for the timsTOF Ultra 2 could likely be ameliorated through the design of a higher capacity tims cartridge. On the part of the user, the method may be modified to reduce the ion accumulation time for higher loads. Pairwise analysis of protein intensities between the Orbitrap Astral and timsTOF Ultra 2 for each sample amount confirmed a generally similar response for the two instruments, with Pearson’s correlation coefficients greater than 0.8 for all sample amounts except at 10 ng where the timsTOF Ultra 2 began to saturate (Figure 3).

**Figure 2.**
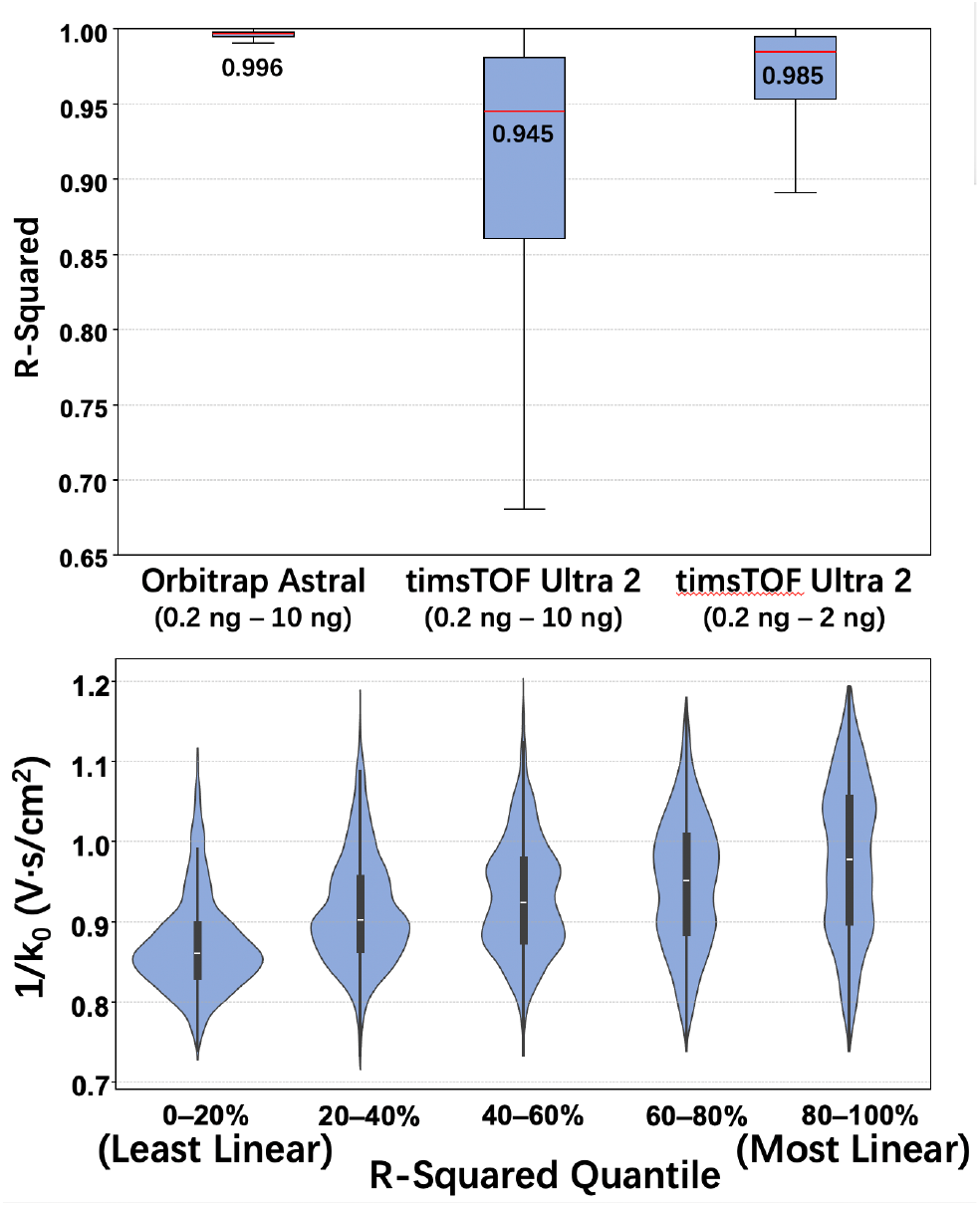
Linearity of reported peptide abundance values across sample amounts. (A) Distribution of R-squared values. (B) Ion mobility vs linearity for peptides analyzed using the timsTOF Ultra 2 mass spectrometer.

**Figure 3.**
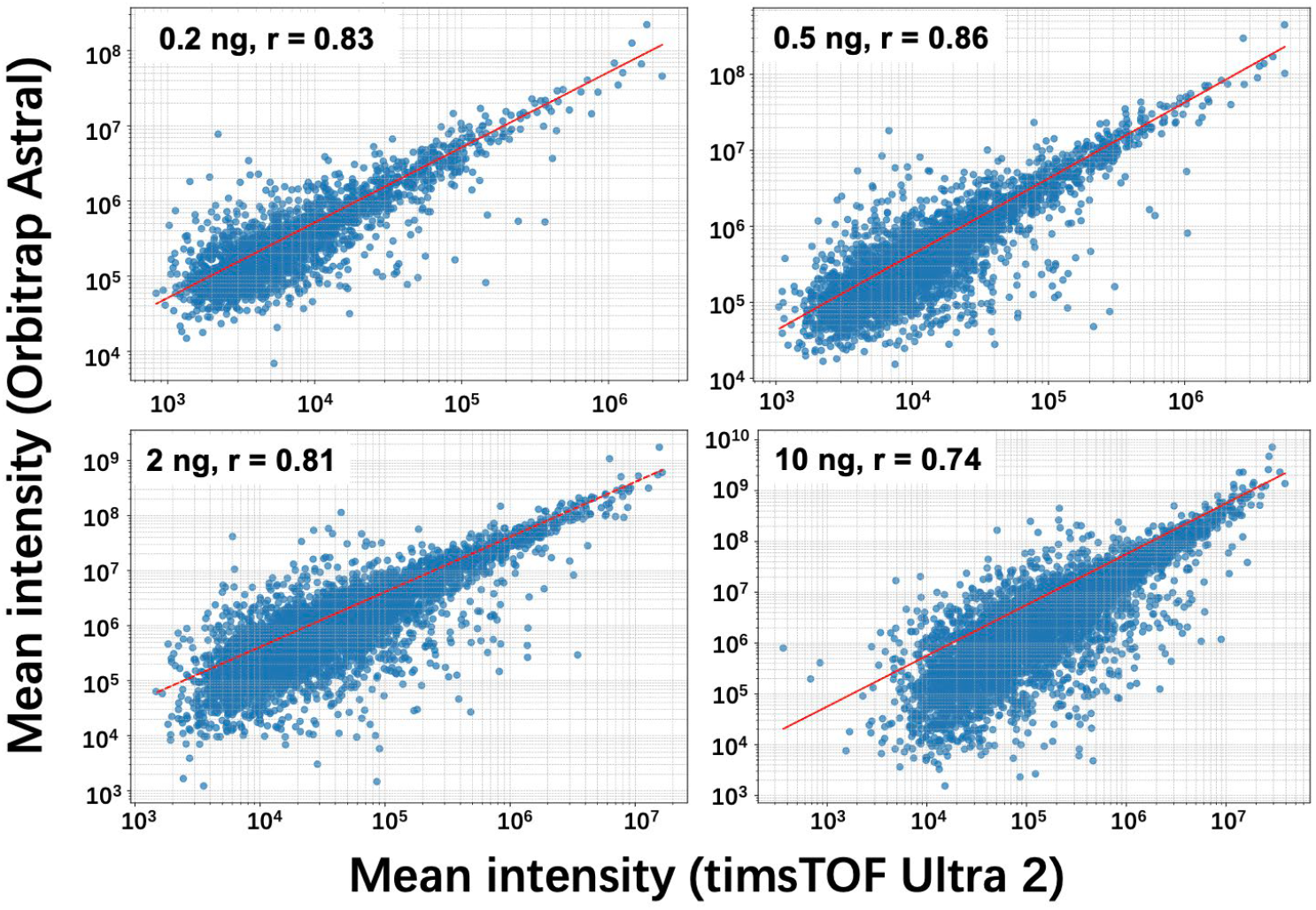
Pairwise correlation of LFQ intensities between the Orbitrap Astral and the timsTOF Ultra 2 for 0.2, 0.5, 2 and 10 ng sample inputs.

### Rapid proteome profiling drives down the cost per sample with minimal impact on proteome depth or reproducibility

While the above analyses point to significant potential cost savings if only low-nanogram samples are prepared and analyzed regardless of sample availability, higher throughput analyses also directly lower the cost per sample since LC and MS instrumentation are primary drivers of cost as detailed above. To assess the impact of gradient duration on proteome depth and reproducibility, we evaluated active gradients from 5 to 60 min on both instruments for 2-ng HeLa digest samples.

The same MS acquisition settings were used as for the above analyses. We were surprised to find that gradient length had a minimal impact on proteome coverage for these rapid, sensitive instruments, as shown in Figure 4A. Indeed, coverage was greatest for the timsTOF Ultra 2 with a 10-min gradient, and only 11% higher than for the 5-min gradient. Similarly, proteome coverage was virtually flat for the Orbitrap Astral from 20–60 min, varying by just 3% within that range and by just 17% across all gradient lengths. We note that the overall cycle time was longer than the gradient lengths shown here, but strategies such as multi-column nanoLC, demonstrated below, can minimize or eliminate the gap between active gradient and overall cycle time once a gradient is selected. Additionally, median CVs of around 10% are achieved across the fastest gradients for both instruments. The identified proteins were also strongly overlapping for both instruments, as shown for 5- and 20-min gradients in Figure 5. Given the minimal added (or even decreased) proteome depth accompanying longer gradients, and the significant reduction in cost per sample that can be achieved with faster cycle times, 5–10 min analyses (144–288 samples per day with 100% duty cycle) should become the norm for global proteome profiling with the current generation of mass spectrometers. The incredible achievable depth with minimal sample input and analysis time also makes these modern mass spectrometers far less expensive than prior generations on a per-sample basis, helping the $10 proteome become a reality.

**Figure 4.**
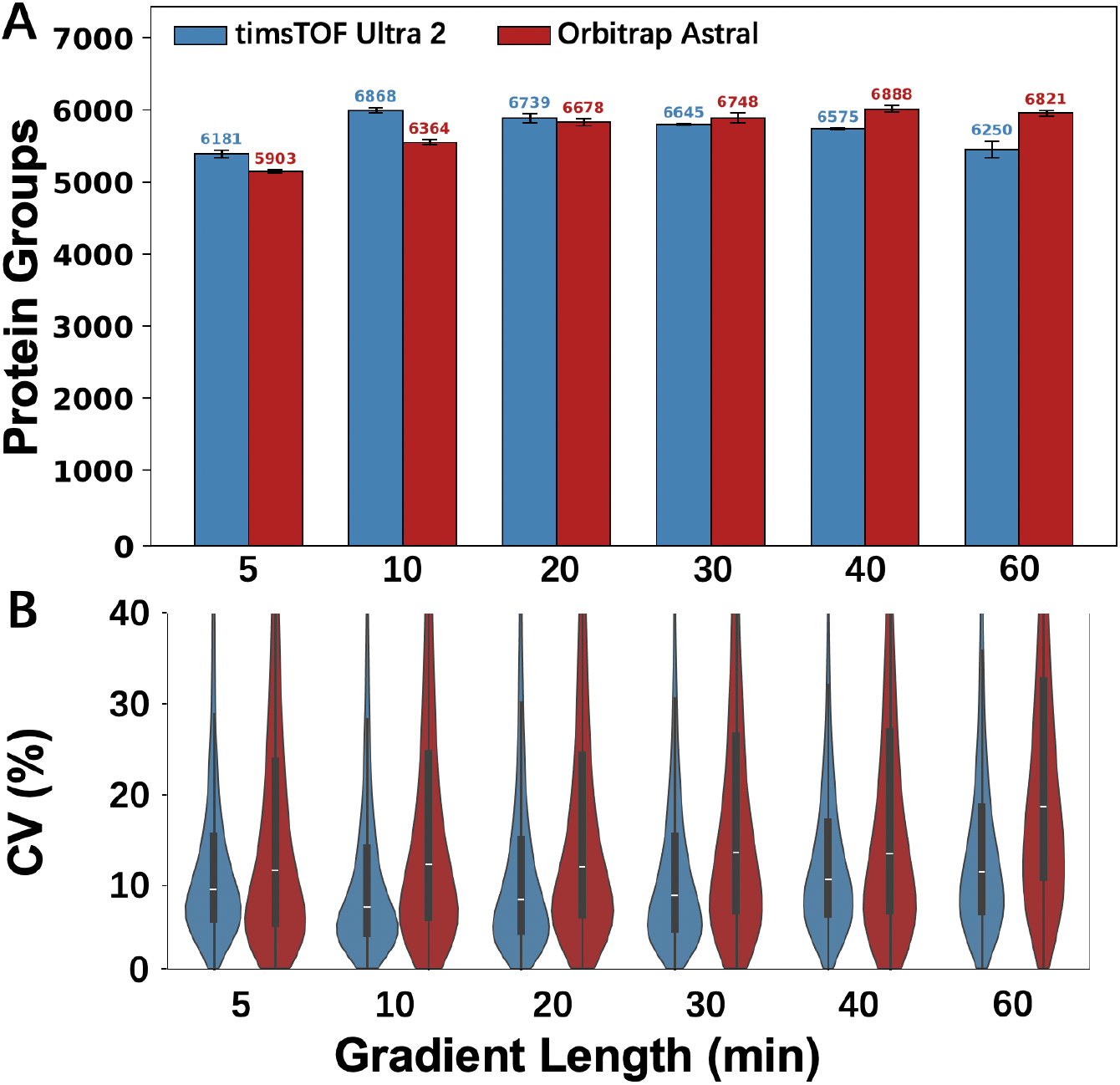
Varied gradient length with a fixed sample input of 2 ng. (A) Proteome coverage. (B) Coefficients of variation.

**Figure 5.**
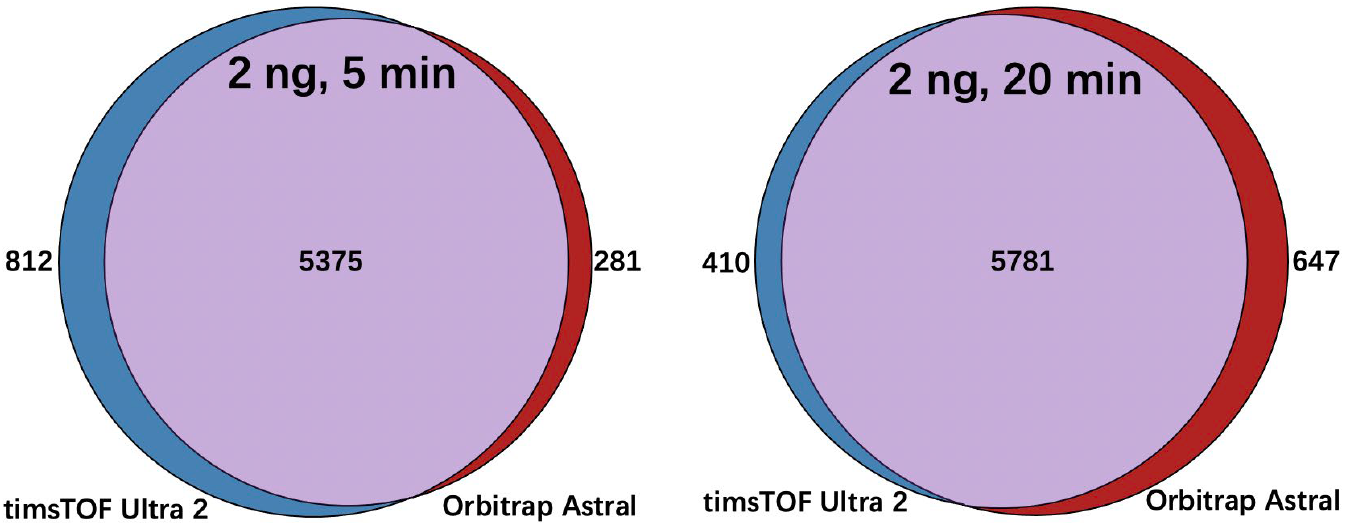
Venn diagrams showing overlap of identified proteins measured with the timsTOF Ultra 2 and the Orbitrap Astral with a 5-min gradient (left) and a 20-min gradient (right).

### Implementation of a 2-column nanoLC system operating at 126 samples per day

The above experiments had some limitations that we sought to address in a follow-up study. First, we did not use the FAIMS Pro interface on the Orbitrap Astral, although it is commonly employed and recommended for low-input analyses. We used nanoLC columns with separate emitters, and we have since found that integrated emitters can produce significantly narrower peaks that further improve performance. For the above study, we employed 30-µm-i.d. columns operated at just 70 nL/min, which is far lower than the flow rate used by most researchers in the field. Finally, according to our calculations, achieving a $10 proteome requires ∼10-min total analysis times or ∼144 samples per day with 80% up time, and our overall cycle time was longer than the reported active gradients. To address many of these limitations, we performed a follow-on experiment on the Orbitrap Astral using a 2-column nanoLC system that had a cycle time of 11.4 min (126 SPD), which is close to the requirement to achieve a 10-min proteome. The duty cycle was nearly 100% for this system, and all experiments (6 technical replicates each for 9 sample amounts) were completed in <10.5 h. One of the 100-ng replicates failed, but all others provided highly consistent results. The FAIMS Pro interface was used for this experiment, and columns with integrated nanoelectrospray emitters were used. The column bore was increased to 50 µm and the flow rate increased to 100 nL/min, which may result in somewhat lower ionization efficiency but is closer to the range of commercial offerings.

We also extended the sample input range to 100 ng to confirm our hypothesis that saturation of proteome coverage occurs with sub-100-ng loadings. This follow-on study thus more fully demonstrates the analytical aspects of the $10 proteome, achieving remarkably deep and reproducible proteome coverage across a range of sample inputs.

While the analytical setup was highly modified relative to the above experiments, proteome coverage and reproducibility as measured by CVs were quite similar. As expected, we found that proteome coverage plateaued, only increasing by <2% above 10 ng as reported by Spectronaut, and by <7% as reported by DIA-NN (Figure 6A). At the lowest inputs, Spectronaut identified nearly 6000 protein groups, while coverage reported by DIA-NN was somewhat lower. Median CVs were impressively less than 10% for all sample inputs >1 ng (Figure 6B). The DIA-NN searches were performed using a spectral library comprising the five 100-ng analyses, and the Spectronaut search was performed in DirectDIA+. All files were analyzed in a batch for both searches. To ensure that the results were not due to spurious assignments by the software when all files are analyzed together, we also analyzed the six lowest input (0.2 ng) samples with just a single 10 ng donor file. As shown in Fig. S1, reported proteome coverage was only modestly lower for Spectronaut using this approach (5,569 vs 5,976 protein groups) and unchanged for DIA-NN, which adds confidence to the findings.

**Figure 6.**
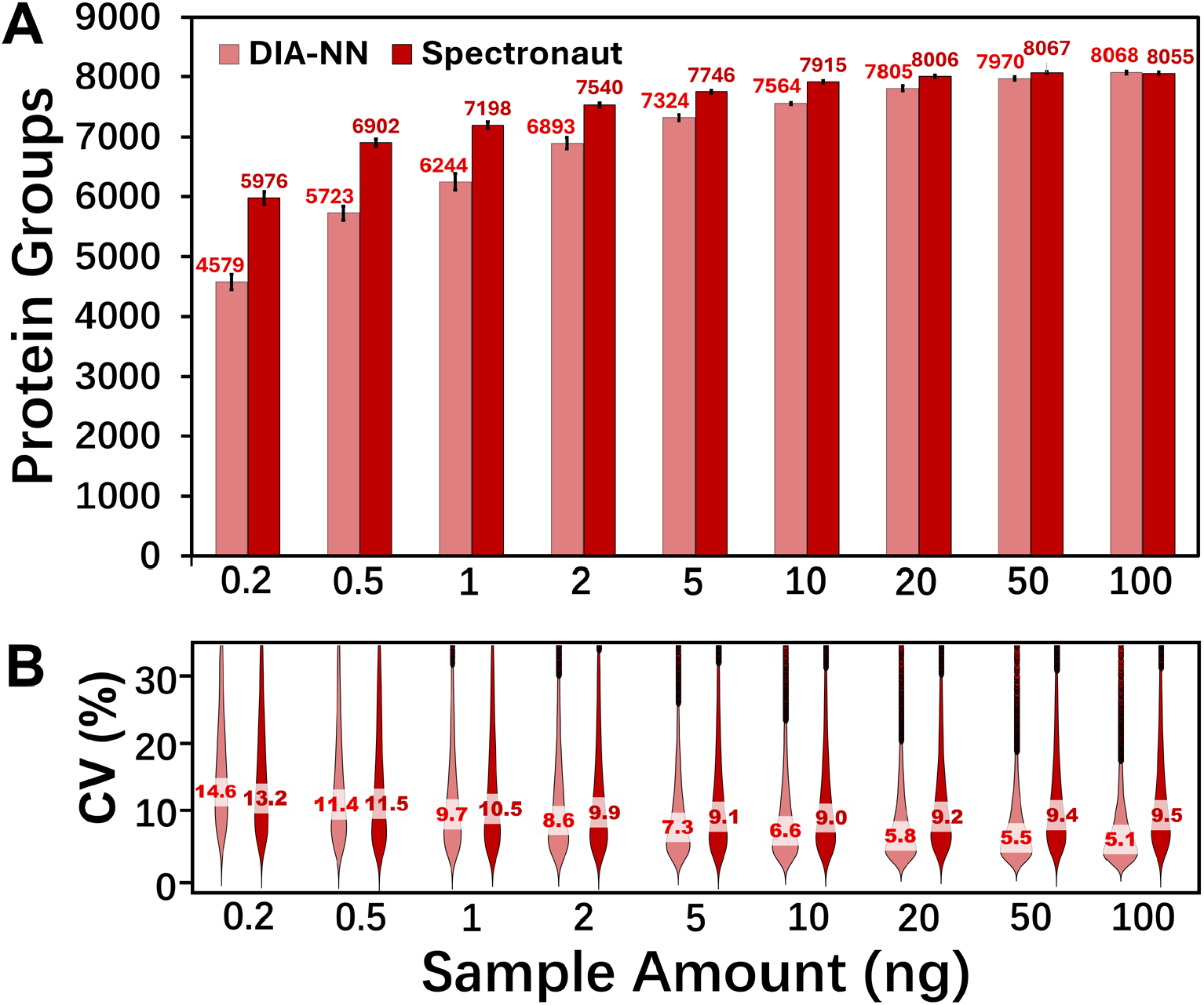
Analysis of sample inputs from 0.2 to 100 ng using a 2-column nanoLC system operated at 126 samples per day (11.4 min cycle time) and the Orbitrap Astral mass spectrometer. (A) Proteome coverage reported by DIA-NN and Spectronaut. (B) Coefficients of variation.

## Conclusion

This study demonstrates that in-depth and reproducible global proteome profiling is now achievable for approximately $10 per sample using current-generation mass spectrometers, specifically the Thermo Fisher Orbitrap Astral and Bruker timsTOF Ultra 2, in combination with simplified, ultra-low-cost sample preparation and standardized data acquisition strategies. By systematically evaluating a wide range of input amounts, gradient durations, and analytical configurations, we show that proteome coverage saturates in the low-nanogram range, and that fixed, non-optimized methods can deliver robust and quantitative results across diverse conditions. The implementation of a multi-column nanoLC system with ∼100% duty cycle pushes throughput to 126 samples per day, while automated data analysis pipelines like MSConnect (32) dramatically reduce labor and expertise barriers. Collectively, these innovations transform proteomics from an expensive and labor-intensive practice into a scalable, affordable platform ready for large-scale biological and clinical studies. Crucially, the $10 proteome paradigm not only democratizes access to proteomics but also positions it to more fully join genomics as a routine, high-throughput pillar of multiomics research. As newer instruments and informatics tools continue to emerge, we anticipate even greater reductions in cost and improvements in depth, reproducibility, and usability. These advances will enable the proteomics community to embrace a new era of standardized, accessible, and impactful protein measurement at scale.

## Methods

### Liquid Chromatography

For experiments corresponding to Figures 1–5, an Ultimate 3000 RSLCnano LC system (Thermo Fisher, Waltham, MA) was operated at a flow rate of 680 nL/min.

The flow was split such that mobile phase passed through the analytical column at ∼70 nL/min, while the remaining flow served as the loading pump for the trap-and-elute sample injection as described in Liang et al. (28). The analytical column was 20 cm long, 30 µm inner diameter (i.d.) and was packed in house with 1.9 µm Dr. Maisch C18 media (Part No. r119.aq.0001, Ammerbuch, Germany). All mobile phase solvents and wash solutions were purchased from Honeywell (Charlotte, NC). Pierce HeLa peptide digest and formic acid ampules were purchased from Thermo Fisher. A 10-port Nanovolume valve (Part No. C72MFSX-4670D) was purchased from VICI (Houston, TX).

The 20-min gradients from solvent A (0.1% formic acid in water) to solvent B (0.1% formic acid in acetonitrile) were as follows: After 1 µL of sample was loaded onto the trap column, the gradient was increased from 1% B to 5% B over 3 minutes, then to 25% B over 20 minutes, to 45% B over 4 minutes, to 80% B over 2 minutes for washing, where it was held for 5 minutes before decreasing to 25% B over 3 minutes and then washing at 80% B for another 5 minutes. The %B was then decreased to 1% where it remained for 11 min to recondition the column. For other gradient lengths, the active gradient from 5–25% B was adjusted while holding other portions of the gradient the same.

For data corresponding to Figure 6 when a 2-column system was used, the LC system architecture matched that described by Xie et al. (52). The LC columns were purchased from MicrOmics Technologies, LLC (Spanish Fork, UT). The lengths remained 20 cm, but the bore was 50 µm and the emitter was integrated with the column. The flow rate was ∼100 nL/min. The total cycle time was 11.4 min, corresponding to 126 samples per day, and the duty cycle (active gradient/cycle time) was close to 100%.

### Mass spectrometry

The data acquisition method for all samples analyzed on the Bruker timsTOF Ultra 2 mass spectrometer was as follows: The LC system was interfaced with a home-etched 10 μm emitter inserted into a CaptiveSpray ESI source, with 1500 V electrospray potential. Full MS data were acquired in the range of m/z 100–1700 and 1.45–0.64 1/K0 [V·s/cm2] in diaPASEF mode. Fifteen DIA windows spanned 400–1000 m/z and were acquired with trapping and ramp times of 150 ms (100% duty cycle). Specific window placement is provided in Figure S2. The estimated cycle time was 0.94 s. The collision energy was ramped as a function of increasing mobility starting from 20 eV at 0.6 1/k_0_ to 63 eV at 1.6 1/k_0_. No denoising mode was applied for diaPASEF data reduction.

With the exception of the data collected with the 2-column nanoLC system corresponding to Figure 6, other samples that were acquired on the Orbitrap Astral mass spectrometer used the following method: A single MS1 scan using the orbitrap detector at 240,000 resolution (at 200 m/z), with the precursor range of m/z 380–980, RF lens setting of 45%, maximum ion injection time set to 100 ms, and the automatic gain control (AGC) target set to 500%. MS2 spectra were simultaneously acquired with the Astral detector, with the following settings: 99 scan events with 4 Da isolation window, 25% HCD collision energy, 45% RF lens, 800% AGC target, 10 ms maximum injection time, and a precursor mass range of 400–800 m/z with window placement optimization. The LC system, column and emitter was the same as for the experiments performed on Bruker timsTOF instrument. The Nanospray Flex ion source was used with an electrospray voltage set to 2200 V. The ion transfer tube temperature was set to 200 °C.

For data acquired using the dual-column nanoLC system (corresponding to Figure 6), the Orbitrap Astral equipped with the FAIMS Pro interface was used. The compensation voltage on the FAIMS interface was fixed at –45 V. The carrier gas flow was 3.6 L/min. Other settings were as described above except for the following settings. MS1 scan range was set to m/z 400-800. The isolation window and injection time was set to 8 Da and 12 ms for sub-2 ng samples, while 4 Da and 6 ms was used for higher-input samples with window placement optimization set to off.

### Data Analysis

All raw files (Bruker .d and Thermo .raw) were analyzed directly with DIA-NN 2.0.2 (53). We followed GitHub page recommendation for DIA-NN: For Bruker timsTOF Ultra data, both mass accuracy and MS1 accuracy were set to 15. For Orbitrap Astral data, mass accuracy was set to 10, and MS1 accuracy was set to 4. Trypsin/P protease specificity allowed up to 1 missed cleavage. Carbamidomethylation of cysteines was a fixed modification, N-term M excision, and precursor charge range of 1+ to 7+. Other precursor and fragment parameters were set to match the MS acquisition methods. All postprocessing was performed using python.

Data presented in Figure 6 were identified and quantified using DIA-NN (v2.0) and Spectronaut (v19.9), with the human UniProt protein database downloaded on July 16, 2025. For the DIA-NN analysis, a custom experimental spectral library was used. This library was constructed from five 100 ng HeLa digest standard runs acquired on the same instrument, using DIA-NN’s library-free mode with deep learning-based identification enabled. Trypsin/P was specified as the digestion enzyme, allowing up to two missed cleavages. Up to five variable modifications were considered, including N-terminal methionine excision, carbamidomethylation (fixed), methionine oxidation, and N-terminal acetylation. The search was performed with a peptide length range of 7–52 amino acids, precursor charge states of 1–4, a precursor m/z range of 400–800, and a fragment ion m/z range of 150–2000. MS1 and MS2 mass accuracies were set to 0 ppm, and precursor FDR was controlled at 1%. Cross-validated neural networks were used for scoring, and cross-run normalization was disabled. The same parameters were applied in the subsequent library-based search, where all raw files acquired on the Orbitrap Astral were analyzed together in a single DIA-NN project.

For the Spectronaut analysis, the directDIA+ workflow was used to analyze all raw files together in a single project. Trypsin/Lys-C was specified as the digestion enzyme, with a peptide length range of 7–52 amino acids and up to two missed cleavages. A maximum of five modifications was allowed per peptide, including fixed carbamidomethylation and variable modifications such as protein N-terminal acetylation and methionine oxidation. The false discovery rate for PSMs, peptides, and protein groups was controlled at 1%. Quantification was performed at the MS1 level, and cross-run normalization was disabled.

Data presented in the Supplementary Information were analyzed using the same workflow described above for DIA-NN and Spectronaut, except that the project consisted of six 200 pg HeLa digest standard runs and one 10 ng HeLa digest standard run, which were processed together in a single project.

## Data Availability

The mass spectrometry proteomic data have been deposited to the ProteomeXchange Consortium via the PRIDE partner repository (54) with the dataset identifier PXD066701.

## Acknowledgments

Research reported in this publication was supported by the National Institute of General Medical Sciences of the National Institutes of Health (NIH) under award number R01GM085232, and by the NIH NCI under award numbers R01CA279074 and U01CA271410 (CPTAC). The content is solely the responsibility of the authors and does not necessarily represent the official views of the National Institutes of Health.

