## Supplementary Figures for "The $10 proteome: low-cost, deep and quantitative proteome profiling of limited sample amounts using the Orbitrap Astral and timsTOF Ultra 2 mass spectrometers"

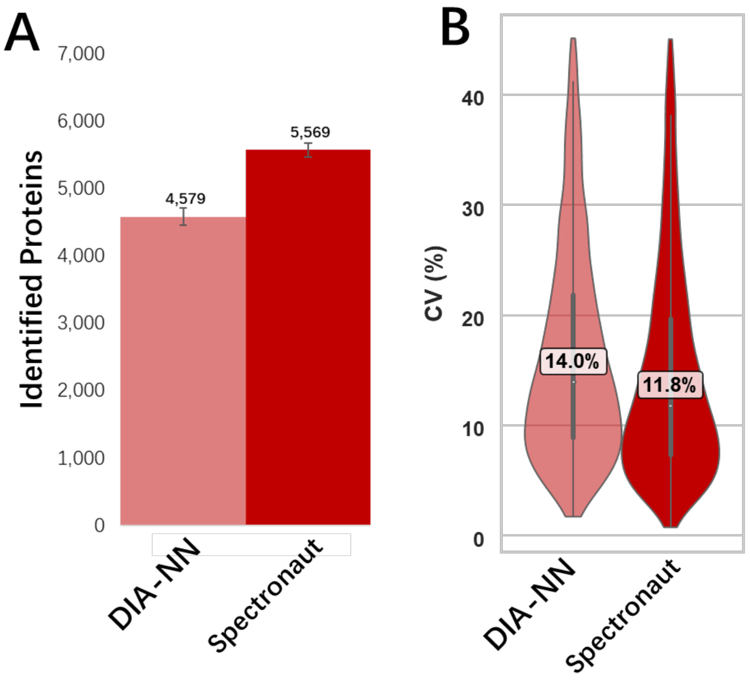


Figure S1. Proteome coverage (A) and coefficients of variation (B) for six 200-pg technical replicates analyzed together with a single 10-ng donor file.

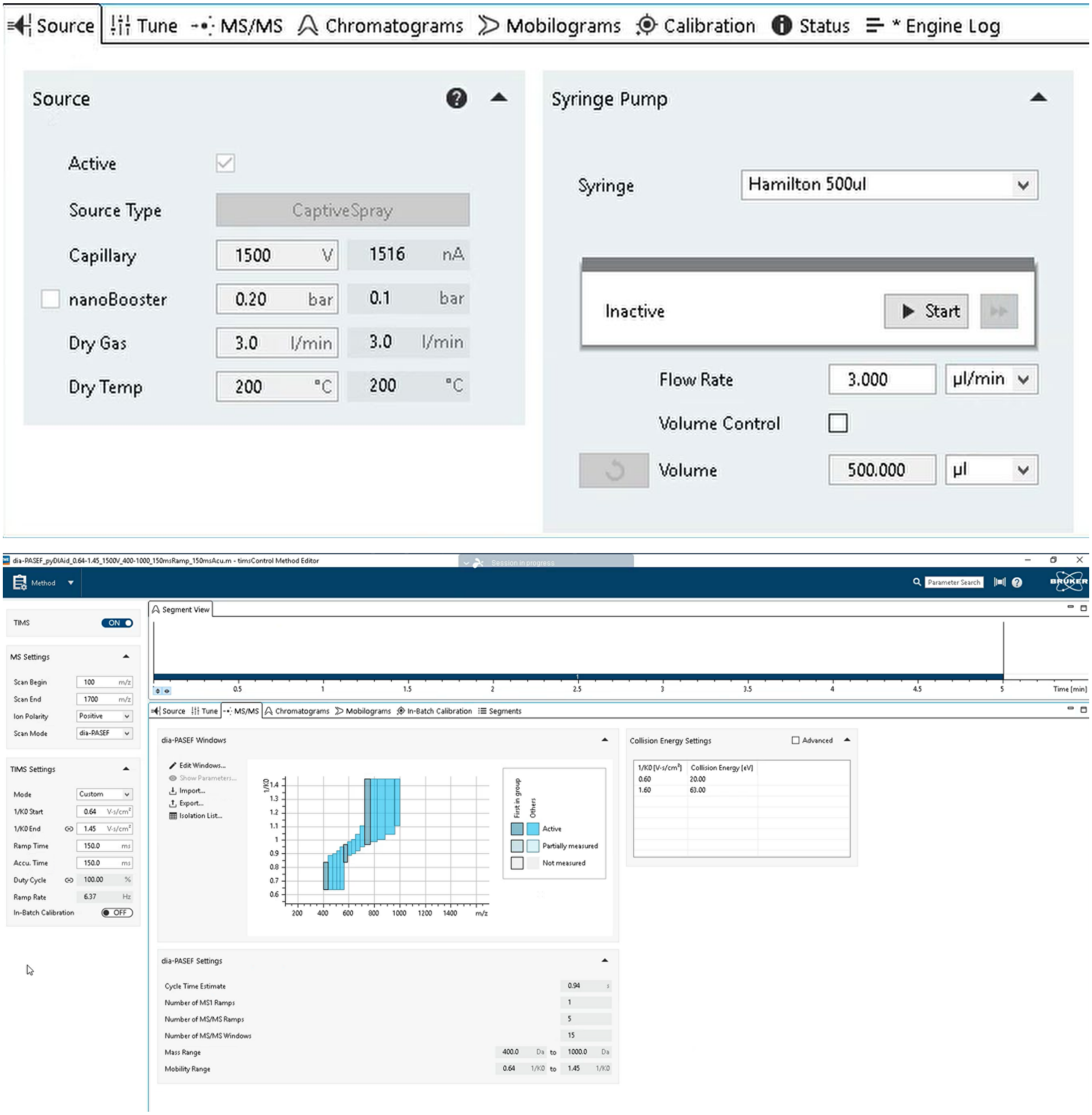


Figure S2. Method used for data acquisition with the timsTOF Ultra 2 mass spectrometer.
